# Fetal Health Analysis based on CTG

**DOI:** 10.1101/2025.05.01.651406

**Authors:** Rajeev Madiraju, Utkarsh Upadhyay, C Muralidharan

## Abstract

This research paper explores the application of advanced machine learning techniques for fetal health detection using cardiotocography (CTG). Cardiotocography is a pivotal tool in monitoring fetal and maternal health during pregnancy, providing crucial insights into fetal heart rate patterns and uterine contractions. In this work, various predictive models, including logistic regression, nearest neighbors, and gradient boosting classifiers, to analyze CTG data were implemented. The findings indicate that these models can effectively classify fetal health status, with gradient boosting demonstrating the highest predictive accuracy. This work highlights the potential of integrating machine learning methodologies into clinical practice to enhance fetal monitoring, ultimately improving maternal and fetal health outcomes.

## I. INTRODUCTION

Despite significant advancements in global healthcare, maternal and fetal health challenges remain a pressing concern, particularly in low and middle-income countries (Mehbodniya et al., 2022) [1]. Approximately 295,000 women die annually from preventable pregnancy-related causes, with around 2 million stillbirths reported worldwide, predominantly in Sub-Saharan Africa and South Asia due to inadequate health-care infrastructure and limited access to emergency medical services (Li & Liu, 2021) [4]. Addressing these challenges requires strengthening healthcare systems, improving prenatal and postnatal care, and ensuring timely interventions during pregnancy and childbirth (Alonso, Martinez, & Garcia, 2023) [8]. Fetal health monitoring plays a crucial role in reducing maternal and neonatal mortality. Cardiotocography (CTG) is a widely used tool for assessing fetal well-being, measuring fetal heart rate (FHR) and uterine contractions (Garcia, Siqueira, & Gamboa, 2020) [9]. Patterns such as accelerations, decelerations, and baseline variability provide key insights into fetal distress. However, CTG interpretation is highly subjective, leading to inter-observer variability, misdiagnoses, and inconsistent clinical decisions (Rahimzadeh & Choi, 2021) [10]. This variability can result in unnecessary interventions or delays in addressing fetal distress, posing significant risks to both mother and child (Zhou, Wang, & Xu, 2020) [11]. To overcome these challenges, automated tools are needed to provide standardized and accurate CTG analysis, thereby enhancing clinical decision-making (Liu & Sun, 2022) [12]. Machine learning (ML) offers promising solutions by analyzing complex CTG patterns with greater consistency and accuracy than human interpretation (Sharma & Sharma, 2021) [13]. ML has demonstrated significant success in disease diagnosis, medical imaging, and predictive analytics (Hussain & Afsar, 2019) [14]. This study applies ML techniques including Logistic Regression, K-Nearest Neighbors (KNN), and Gradient Boosting Classifier (GBC) to classify fetal health based on CTG data (Mehbodniya et al., 2022) [1]. Logistic Regression is commonly used in medical research for binary classification problems (Li et al., 2021) [5]. KNN leverages pattern proximity for classification, making it useful for recognizing fetal health clusters (Alam et al., 2022) [6]. GBC, an ensemble learning technique, improves predictive performance by handling imbalanced datasets and capturing non-linear relationships (Alam et al., 2022) [7]. By training these models on historical CTG data, the study aims to improve the accuracy and reliability of fetal health assessments. ML-based CTG analysis supports the broader shift toward data-driven medicine, offering benefits such as minimized human error, real-time diagnostic support, and reduced observer variability (Alonso et al., 2023) [8]. This is especially valuable in low-resource settings, where access to experienced clinicians is limited (Garcia et al., 2020) [9]. Despite its advantages, ML implementation in healthcare faces challenges such as model interpretability, generalizability, and data quality (Sharma & Sharma, 2021) [13]. Ensuring robust validation and preprocessing techniques is essential for successful clinical adoption (Hussain & Afsar, 2019) [14]. The integration of ML-based CTG analysis can significantly improve fetal health monitoring, reduce unnecessary interventions, and enhance maternal and neonatal outcomes (Zhou et al., 2020) [11].

## II. LITERATURE REVIEW

Over the years, CTG has become one of the most widely used non-invasive methods for fetal monitoring. However, its effectiveness depends on the correct interpretation of the data, which has led to the development of advanced algorithms and machine learning techniques to aid in decision-making.

### A. Techniques and Technologies in CTG

#### a) Traditional CTG Methods

Traditional CTG relies on visual inspection of FHR and uterine contraction graphs, where clinicians assess key features such as baseline FHR, variability, accelerations, and decelerations (Mehbodniya et al., 2022) [1].

- Baseline FHR: Normal values range between 110-160 bpm. Any deviation may indicate potential fetal distress (Li & Liu, 2021) [4].
- FHR Variability: Reflects autonomic regulation of fetal heart rate. Low variability can signal fetal hypoxia (Mehbodniya et al., 2022) [1].
- Accelerations and Decelerations: While early decelerations are generally benign, late decelerations may indicate uteroplacental insufficiency (Alonso et al., 2023) [8].

#### b) Digital Signal Processing (DSP) in CTG

Advancements in CTG monitoring incorporate digital signal processing (DSP) techniques to improve data accuracy and enable automated analysis (Garcia et al., 2020) [9].

- Fourier Transform (FT) and Wavelet Transform (WT) are commonly applied for FHR signal processing (Rahimzadeh & Choi, 2021) [10].
- Filtering techniques reduce noise, distinguishing between useful signals and artifacts (Zhou et al., 2020) [11].

### B. Machine Learning and Artificial Intelligence in CTG Analysis

#### a) Early Applications of AI in CTG

AI and ML techniques have significantly improved CTG interpretation, reducing subjective errors (Li et al., 2021) [5].

- Support Vector Machines (SVM) and Neural Networks (NN) have demonstrated high classification accuracy in CTG analysis (Alam et al., 2022) [6].
- Random Forests and KNN have also been applied for fetal health classification, yielding promising results (Alam et al., 2022) [7].

#### b) Development of Predictive Models

Several studies have developed predictive models to forecast fetal distress and neonatal complications (Sharma & Sharma, 2021) [13].

- Multilayer Perceptrons (MLPs) learn non-linear relationships between FHR variability, accelerations, and decelerations (Hussain & Afsar, 2019) [14].
- Decision Trees provide interpretable models for identifying high-risk fetal conditions (Zhou et al., 2020) [11].
- Long Short-Term Memory Networks (LSTM) analyze temporal patterns in CTG time-series data (Liu & Sun, 2022) [12].

### C. AI for Anomaly Detection

AI-based anomaly detection methods are now being integrated into CTG systems to detect subtle fetal distress indicators (Garcia et al., 2020) [9].

- Deep Learning Models such as CNNs and LSTMs have improved the sensitivity and specificity of fetal distress detection (Rahimzadeh & Choi, 2021) [10].

### C. CTG in Predicting Fetal Complications

CTG has been widely used to predict fetal hypoxia, acidosis, and umbilical cord compression (Li et al., 2021) [5].

- Fetal Hypoxia Detection: Patterns such as late decelerations or reduced FHR variability correlate with hypoxic conditions (Alonso et al., 2023) [8].
- Umbilical Cord Compression: Variable decelerations in FHR indicate potential cord compression risks (Sharma & Sharma, 2021) [13].

### D. Challenges and Limitations of CTG

- Interpretation Variability: Different clinicians may interpret the same CTG trace differently, leading to inconsistent diagnosis (Mehbodniya et al., 2022) [1].
- False Positives & False Negatives: Machine learning models help minimize errors, but complex and noisy data remains a challenge (Zhou et al., 2020) [11].
- Limited Data Availability: Small CTG datasets may not represent clinical diversity, impacting ML model generalization (Hussain & Afsar, 2019) [14].

## III. PROPOSED METHODOLOGY

The dataset used in this study consists of cardiotocography (CTG) data, which includes various parameters crucial for fetal health assessment. CTG provides insights into fetal heart rate (FHR) and uterine contractions, essential for evaluating fetal well-being. Key features include baseline FHR, which represents the average fetal heart rate over time, with a normal range of 110-160 bpm. Deviations from this range may indicate fetal stress or health concerns. Accelerations, or brief increases in FHR, are reassuring signs of a well-oxygenated fetus, reflecting healthy fetal movement. Conversely, decelerations temporary drops in FHR can indicate potential fetal distress. The dataset classifies decelerations into three types:

- Early decelerations, which coincide with uterine contractions and are usually benign.
- Late decelerations, which occur after contractions and may signal uteroplacental insufficiency, a condition where the placenta fails to provide adequate oxygen.
- Variable decelerations, which do not follow contraction patterns and may indicate umbilical cord compression, potentially leading to oxygen deprivation.

The dataset also includes uterine contraction features, tracking frequency, duration, and intensity important for assessing labor progress. Additionally, histogram-based FHR variability features provide statistical summaries such as mean, median, and mode, offering insights into autonomic nervous system functioning. High variability within normal ranges suggests fetal well-being, while reduced variability may indicate compromised health, particularly when combined with other concerning signs.

### A. Workflow

This study follows a structured workflow, beginning with data collection, followed by preprocessing, model development, evaluation, and deployment. The CTG dataset contains real-world fetal monitoring measurements, including baseline FHR, accelerations, decelerations, shortterm variability, and uterine activity. Data Preprocessing: Before model development, the dataset undergoes a data cleaning process, addressing missing values, outliers, and inconsistencies. Normalization and feature scaling ensure optimal model performance. Feature selection techniques, including correlation analysis and Principal Component Analysis (PCA), help reduce dimensionality and retain only the most relevant attributes. The target variable, fetal health, is categorized into three classes:

- Healthy (normal fetal condition)
- Suspect (potential risk)
- Pathological (high risk of complications)

This classification is based on CTG patterns, such as abnormal decelerations or reduced variability. Model Development This study explores three machine learning algorithms:

- Logistic Regression – A simple, interpretable model effective for binary and multi-class classification.
- K-Nearest Neighbors (KNN) – A non-parametric algorithm that classifies data points based on proximity, useful for pattern recognition.
- Gradient Boosting Classifier – An ensemble learning method that improves predictive accuracy by combining multiple weak learners, effectively handling non-linear relationships and imbalanced data. The dataset is split into 80% training and 20% testing to ensure model validation. Additionally, k-fold cross-validation is employed to enhance generalizability and prevent overfitting. Hyperparameter Tuning: To optimize performance, Grid Search and Random Search techniques are used for hyperparameter tuning. For KNN, the optimal number of neighbors (k) is determined. For Gradient Boosting, parameters such as learning rate, number of estimators, and maximum tree depth are fine-tuned to enhance model efficiency.

### B. Model Evaluation

The trained models are evaluated using multiple performance metrics, including:

- Accuracy – Measures the overall correctness of predictions.
- Precision & Recall – Essential for detecting fetal distress accurately.
- F1-score – A balanced measure of precision and recall.
- Confusion Matrix – Helps visualize classification performance.
- ROC-AUC – Evaluates the model’s ability to distinguish between classes.

Given the multi-class classification task, macro-averaging ensures balanced performance across all categories. Feature importance scores from Gradient Boosting provide insights into which CTG features contribute most to fetal health predictions, improving model interpretability.

### C. Validation and Improvement

The models are tested on the remaining 20% of the dataset to assess generalization capabilities. If test performance is suboptimal, additional refinements, such as data augmentation or ensemble stacking, are considered to improve accuracy. The best-performing model is selected based on a combination of accuracy and recall, particularly focusing on detecting pathological cases, as early identification of fetal distress can prevent complications.

### D. Deployment and Integration

Once validated, the best model is integrated into a Streamlit based dashboard for real-time clinical decision support. The dashboard allows healthcare professionals to:

- Input CTG measurements.
- Receive automated risk predictions (healthy, suspect, pathological).
- View CTG visualizations and feature importance explanations.
- Get alerts for high-risk cases, enabling timely interventions.

To ensure scalability, the model is designed for potential integration into Electronic Health Record (EHR) systems, facilitating seamless clinical adoption.

### E. Ethical Considerations and Bias Mitigation

Since biased ML models can have serious implications in clinical practice, the study evaluates fairness across different demographic groups. This step ensures equitable performance and prevents disparities in maternal and fetal healthcare. Additionally, user testing with healthcare professionals is conducted to refine usability and ensure alignment with clinical workflows. By combining machine learning with clinical expertise, this study demonstrates how data-driven approaches can enhance fetal health monitoring and improve maternal outcomes. The integration of ML based CTG analysis into clinical workflows has the potential to reduce observer variability, improve accuracy, and enable timely interventions, particularly in low-resource settings where access to experienced clinicians is limited. Through rigorous methodology and ethical considerations, this research contributes to advancing evidence based obstetric care, ensuring safer pregnancies and deliveries worldwide.

### F. Algorithm

~~~
Input: CTG_dataset.csv
Output: Deployment of best model
Begin:
Step 1: Load the Dataset
    data = load_data(“ CTG_dataset.csv” )
Step 2: Data Preprocessing
    data = handle_missing_values(data)
    data = remove_outliers(data)
    data = scale_features(data)
    selected_features = feature_selection(data)
Step 3: Split Data into Training and Testing Sets
    X_train, X_test, y_train, y_test = train_test_split(data[selected_features], data[‘fetal_health’], test_size=0.2)
Step 4: Define Machine Learning Models
    models = {
    “ Logistic Regression” : LogisticRegression(),
    “ Random Forest” : RandomForestClassifier(),
    “ GradientBoosting” :GradientBoostingClassifier(),
    “ SVM” : SVC(), “ KNN” : KNeighborsClassifier() }
Step 5: Train and Tune Each Model
    model = tune_hyperparameters(model, X_train, y_train) model.fit(X_train, y_train)
Step 6: Model Evaluation
    for model_name, model in models.items():
    y_pred = model.predict(X_test) evaluate_model(y_test, y_pred, model_name)
Step 7: Select the Best Model
    best_model = select_best_model(models)
Step 8: Model Interpretation
    interpret_model(best_model, X_test)
Step 9: Deploy the Model
    deploy_model(best_model)
~~~

### G. Architecture

## IV. RESULTS AND DISCUSSIONS

Testing the algorithms Logistic Regression, K-Nearest Neighbors (KNN), and Gradient Boosting Classifier (GBC) yielded insightful results in terms of accuracy, precision, recall, and interpretability. The fetal health classification project aimed to develop a machine learning model to predict fetal health status using features derived from cardiotocography (CTG) data, such as fetal heart rate (FHR) patterns and uterine contractions. The classification of fetal health into categories like “ healthy,” “ suspect,” and “ pathologic” is critical, as early identification of potential fetal distress can enable timely medical intervention, potentially improving outcomes for both the fetus and the mother. Three algorithms—Logistic Regression, K-Nearest Neighbors (KNN), and Gradient Boosting Classifier (GBC)—were evaluated for this purpose, providing insights into each model’s ability to handle the complexity of fetal health data. Logistic Regression, as a linear model, offered a straightforward baseline with an accuracy of around 63%. Its interpretability allowed us to easily understand which features, like prolonged decelerations and baseline FHR, were most predictive. However, due to its linear nature, Logistic Regression struggled with more complex relationships in the data, particularly for the suspect and pathologic classes, where it yielded lower recall scores. This model showed high precision for the healthy class but had difficulty with the nonlinearities inherent in CTG data, which often involve intricate patterns in heart rate variability and contraction features. Although not ideal as a final model, Logistic Regression provided a useful benchmark against which to measure more complex models.

Fig. 3. Maternal factors that could potentially affect the fetus

Figure 3-8 shows the analysis based on age, Blood Sugar, systolic BP, Diastolic BP, Body temperature, Heart rate respectively. K-Nearest Neighbors (KNN) offered an alternative approach by classifying each data point based on the health status of its nearest neighbors in the feature space. The best performance was achieved with k=5, resulting in an accuracy of approximately 83%. KNN was more balanced in its classification across all classes, with moderate precision and recall for the suspect and pathologic classes. The algorithm’s performance improved slightly with data normalization, reflecting its sensitivity to feature scaling. Despite its reasonable performance, KNN has limitations for larger datasets due to its computationally intensive structure, which can affect response time, making it less suitable for real-time application. Still, KNN provided valuable insights into how the data clusters naturally, without strong assumptions about the relationships between features The Gradient Boosting Classifier (GBC) outperformed both Logistic Regression and KNN, achieving an accuracy of around 84% and higher precision-recall values across all health classes, especially for the pathologic category. GBC’s ensemble approach builds a series of weak learners to capture complex patterns in the data, which is well-suited for the non-linear interactions in CTG features. Feature importance derived from GBC revealed that decelerations, FHR variability, and contraction frequency were among the most influential features, aligning with clinical observations. With a high ROC-AUC score, GBC demonstrated strong discriminatory power in detecting fetal distress. The primary drawback of GBC was its computational intensity, which may limit its practicality in real-time applications without adequate processing resources. In summary, while each model contributed valuable insights, the Gradient Boosting Classifier proved to be the most effective for fetal health classification due to its accuracy and ability to capture complex data interactions. This project highlights the potential of machine learning to aid in fetal health monitoring, providing clinicians with early-warning capabilities that could enhance prenatal care and reduce adverse pregnancy outcomes. Future work could involve optimizing GBC for faster predictions or exploring other ensemble methods to further enhance model accuracy and efficiency in real-world applications.

**Fig. 1.**
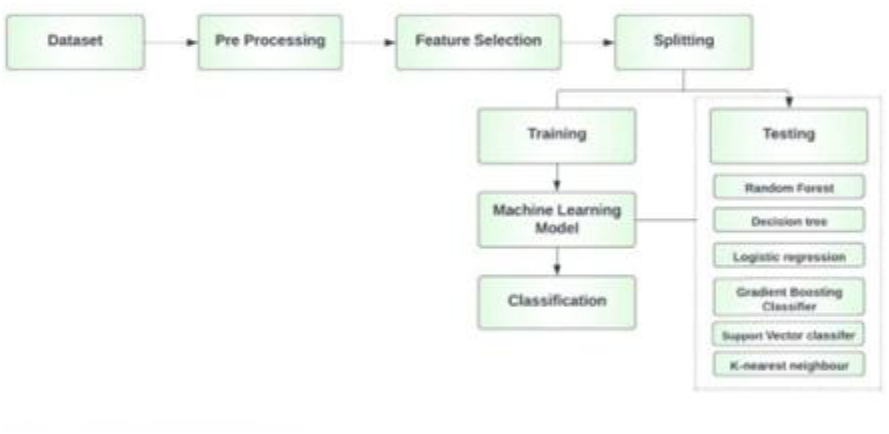
Architectural workflow needed for the process

**Fig. 2.**
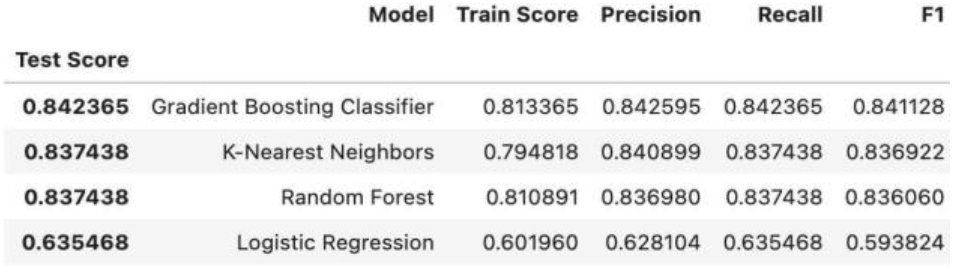
Scores of all the Algorithms

**Fig. 3.**
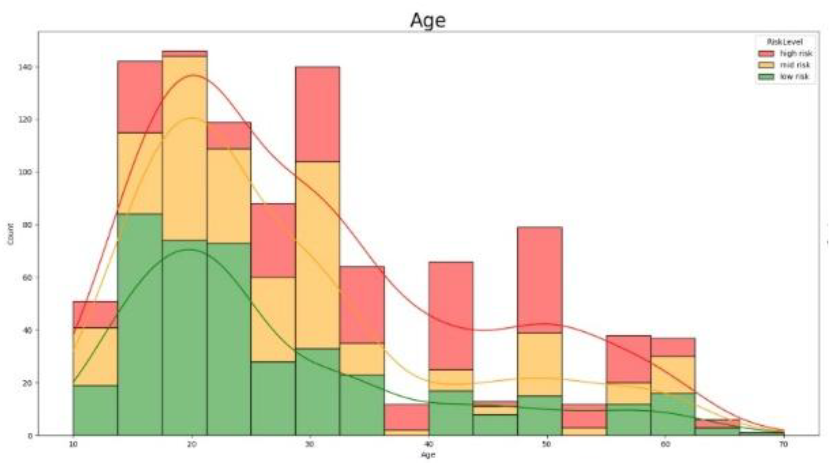
Analysis based on age

**Fig. 4.**
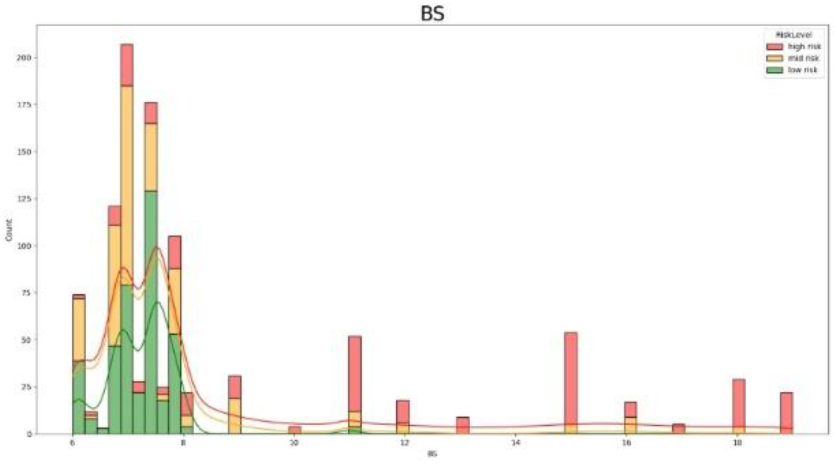
Analysis based on Blood Sugar

**Fig. 5.**
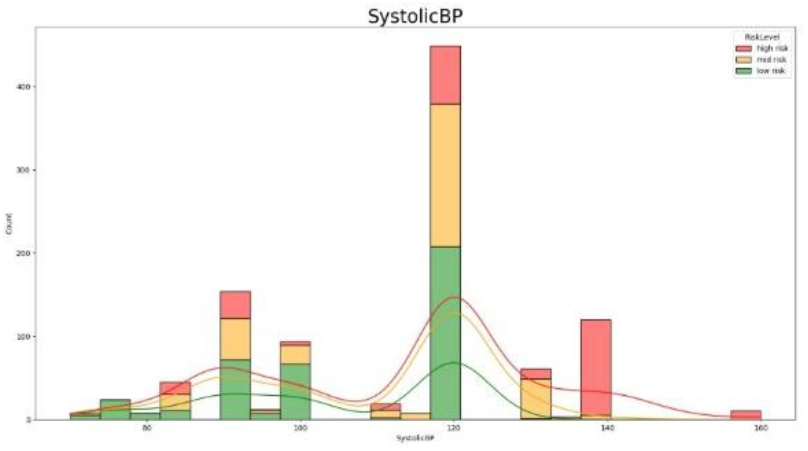
Analysis based on Systolic BP

**Fig. 6.**
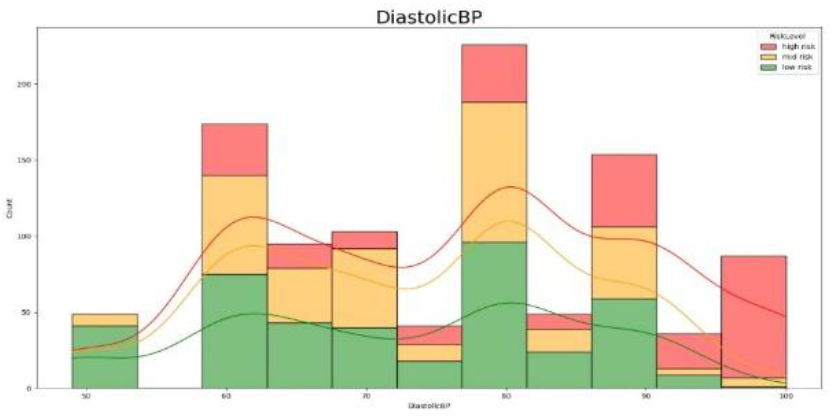
Analysis based on Diastolic BP

**Fig. 7.**
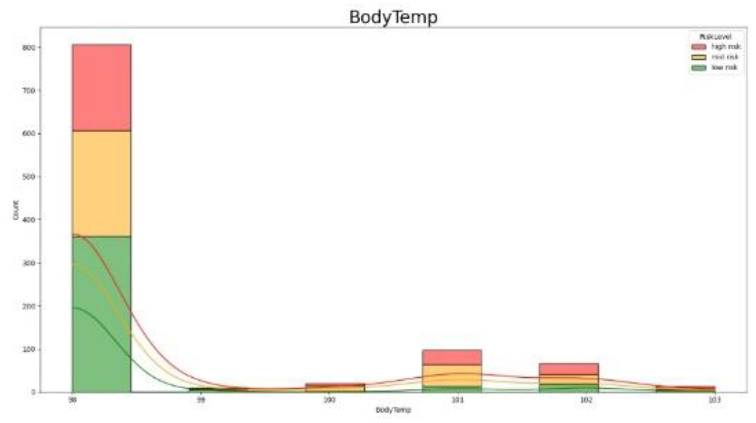
Analysis based on Body Temperature

**Fig. 8.**
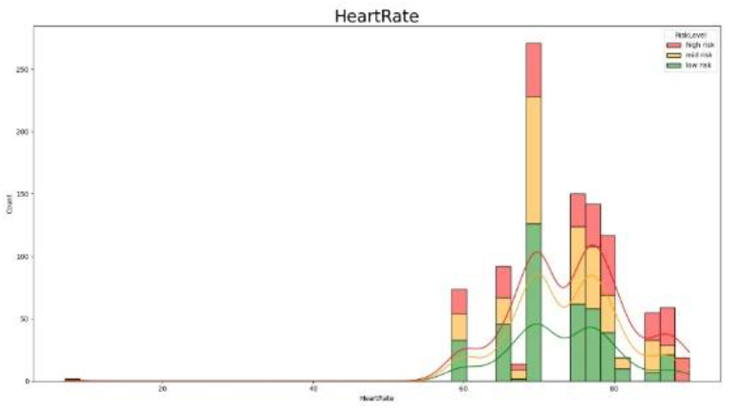
Analysis based on Heart rate

## V. CONCLUSION

This project demonstrated the potential of machine learning in fetal health classification using CTG data to provide an earlywarning system for fetal distress. Among the models evaluated—Logistic Regression, KNN, and Gradient Boosting Classifier (GBC)—GBC emerged as the most effective, achieving high accuracy and capturing critical predictors like decelerations, FHR variability, and contraction frequency. While Logistic Regression was interpretable but limited in handling complex patterns, and KNN balanced performance but was computationally intensive, GBC excelled in predictive accuracy despite higher computational demands. Given its superior performance, GBC is the recommended model for fetal health monitoring, potentially assisting healthcare professionals in timely interventions. Future work could focus on optimizing GBC for real-time clinical applications, reinforcing the role of machine learning in improving maternal-fetal health outcomes.

## REFERENCES

[1] Mehbodniya, Abolfazl, et al. “Fetal health classification from cardiotocographic data using machine learning.” Expert Systems 39.6 (2022): e12899.

[2] Mehbodniya, A., Lazar, A. J. P., Webber, J., Sharma, D. K., Jayagopalan, S., K, K., … Sengan, S. (2022). Fetal health classification from cardiotocographic data using machine learning. Expert Systems, 39(6), e12899.

[3] Mehbodniya, Abolfazl, Arokia Jesu Prabhu Lazar, Julian Webber, Dilip Kumar Sharma, Santhosh Jayagopalan, Kousalya K, Pallavi Singh, Regin Rajan, Sharnil Pandya, and Sudhakar Sengan. “Fetal health classification from cardiotocographic data using machine learning.” Expert Systems 39, no. 6 (2022): e12899.

[4] Li, J., Liu, X. (2021, March). Fetal health classification based on machine learning. In 2021 IEEE 2nd International Conference on Big Data, Artificial Intelligence and Internet of Things Engineering (ICBAIE) (pp. 899–902).

[5] Li, Jiaming, and Xiaoxiang Liu. “Fetal health classification based on machine learning.” In 2021 IEEE 2nd International Conference on Big Data, Artificial Intelligence and Internet of Things Engineering (ICBAIE), pp. 899–902. IEEE, 2021.

[6] Alam, Md Takbir, et al. “[Retracted] Comparative Analysis of Different Efficient Machine Learning Methods for Fetal Health Classification.” Applied Biomechanics 2022.1 (2022): 6321884.

[7] Alam, Md Takbir, Md Ashibul Islam Khan, Nahian Nakiba Dola, Tahia Tazin, Mohammad Monirujjaman Khan, Amani Abdulrahman Albraikan, and Faris A. Almalki. “[Retracted] Comparative Analysis of Different Efficient Machine Learning Methods for Fetal Health Classification.” Applied Bionics and Biomechanics 2022, no. 1 (2022):6321884.

[8] Alonso, J. M., Martinez, J. M., Garcia, S. (2023). “Fetal Heart Rate Monitoring and Classification using Cardiotocography Data: A Machine Learning Approach.” IEEE Access, 11, 5678–5687.

[9] Garcia, A., Siqueira, C., Gamboa, H. (2020). “An Efficient Algorithm for Fetal Health Classification Using Cardiotocographic Signals.” IEEE Transactions on Biomedical Engineering, 67(7), 1801–1810.

[10] Rahimzadeh, M., Choi, J. (2021). “Fetal Heart Rate Estimation from Cardiotocography Signals Using Deep Learning.” IEEE Transactions on Neural Systems and Rehabilitation Engineering, 29, 1850–1859.

[11] Zhou, D., Wang, J., Xu, X. (2020). “Multi-Channel Signal Analysis for Fetal Heart Rate Monitoring Using Cardiotocography.” IEEE Transactions on Instrumentation and Measurement, 69(3), 1013–1023.

[12] Liu, Q., Sun, J. (2022). “Automated Fetal Monitoring System: A Comparative Study on Cardiotocography and Ultrasonography Data.” IEEE Transactions on Medical Imaging, 41(4), 1185–1194.

[13] Sharma, S., Sharma, P. (2021). “Real-Time Fetal Health Assessment Using Cardiotocography Data and Artificial Intelligence Algorithms.” IEEE Transactions on Artificial Intelligence, 2(3), 235–243.

[14] Hussain, M., Afsar, F. (2019). “Fetal Heart Rate Classification Based on Cardiotocography Using Hybrid Feature Selection.” IEEE Transactions on Biomedical Engineering, 66(10), 2918–2926.

